# RAG-17: A Novel siRNA Conjugate Demonstrating Efficacy in Late-Stage Treatment of SOD1^G93A^ ALS mice

**DOI:** 10.1101/2023.11.23.568255

**Authors:** Chunling Duan, Moorim Kang, Kunshan Liu, Zubao Gan, Guanlin Li, Junnan Chen, Ian Schacht, Robert F. Place, Long-Cheng Li

## Abstract

Amyotrophic lateral sclerosis (ALS) is a devastating neurodegenerative disease characterized by rapid progression and high mortality. With genetic mutations, particularly in the SOD1 gene, playing a significant role in ALS pathogenesis, targeted therapies have become a primary focus. This study introduces RD-12500 (RAG-17), a novel siRNA-ACO (Accessory Oligonucleotide) conjugate designed to address the challenges of delivering duplex RNAs to the central nervous system (CNS). RD-12500 exhibits remarkable *in vitro* stability and target specificity with minimal immunostimulation. *In vivo* studies demonstrate its extensive CNS biodistribution, sustained accumulation post-intrathecal administration, and a robust dose-exposure-activity correlation. Notably, RD-12500 significantly reduces cerebrospinal fluid (CSF) SOD1 protein levels, indicating potent SOD1 mRNA and protein knockdown in cynomolgus monkeys. Most notably, our study breaks new ground by demonstrating the effectiveness of RD-12500 in late-stage treatment scenarios. In SOD1^G93A^ ALS mice, post-onset administration of RD-12500 significantly delayed disease progression, improved motor function, and extended survival, marking a significant advancement over other treatments which are typically initiated pre-symptomatically in the same model mice. These findings suggest RD-12500’s potential to provide therapeutic benefits not only to pre-symptomatic but also to post-symptomatic and late-stage SOD1-ALS patients.

## INTRODUCTION

Amyotrophic lateral sclerosis (ALS), a cluster of progressive neurodegenerative conditions, is marked by the debilitating loss of both upper and lower motor neurons. This leads to muscle weakness, paralysis, and ultimately, death within 2-5 years of diagnosis. ALS manifests predominantly as sporadic (90% of cases), while familial forms constitute about 10% of cases) (1, 2).

Over 30 genetic mutations have been identified in association with ALS pathogenesis. Notably, mutations in the SOD1 gene are responsible for approximately 20% of familial ALS cases and 2-3% of all ALS cases (2-4). In Asian populations, SOD1 mutations are the leading cause of familial ALS (5, 6). Mutated SOD1 is believed to contribute to motor neuron degeneration primarily through the toxic aggregation of misfolded proteins (7).

Targeting the reduction of mutant SOD1 protein expression has emerged as a promising therapeutic strategy for SOD1-ALS. In this context, Tofersen, an antisense oligonucleotide (ASO) targeting the 3’-untranslated region (3’UTR) of SOD1 mRNA for RNaseH-mediated degradation, has recently gained accelerated FDA approval for treating SOD1-ALS, following its demonstrated efficacy in lowering plasma neurofilament light chain levels, indicative of neuro-axonal damage (8).

Despite the progress, clinical development of oligonucleotide therapeutics for neurodegenerative diseases has been predominantly limited to single-stranded ASOs. These ASOs, when administered locally (e.g., intrathecally), exhibit significant CNS biodistribution and activity. This has led to the approval of ASO drugs like Tofersen and Nusinersen, with several others in clinical development (9). However, duplex RNA therapies, such as siRNA and saRNA, face substantial delivery challenges for modulating CNS targets, and only one siRNA program reaching Phase I trials for Alzheimer’s disease)(10).

To address these delivery challenges, we have pioneered a novel conjugate – a combination of ASO’s self-delivering properties and the potent knockdown capability of duplex siRNA, which we term smart chemistry aided delivery (SCAD). Our study demonstrates that SOD1 siRNA linked to a non-targeting ASO (Accessory Oligonucleotide, ACO) exhibits strong *in vivo* efficacy in SOD1^G93A^ mice, significantly reducing SOD1 expression and ameliorating disease phenotypes (11).

In the present work, we further characterized the *in vitro* stability, secondary pharmacology of the SOD1 siRNA-ACO conjugate RD-12500 (RAG-17), and its pharmacokinetics and pharmacodynamics in rats and monkeys. Our results reveal compelling therapeutic potential for RD-12500 in SOD1^G93A^ rats. Furthermore, post-onset treatment with RD-12500 in SOD1^G93A^ mice significantly mitigated ALS symptoms and extended survival, underscoring its potential for post-symptomatic treatment in SOD1-ALS patients. This work lays a solid foundation for advancing RAG-17 into clinical trials for the treatment of SOD1-ALS.

## RESULTS

### *In vitro* characterization of RD-12500

#### Serum stability of RD-12500

To evaluate the serum stability of RD-12500 (also referred to as RAG-17), which is a duplex of SOD1 siRNA linked to an accessory oligonucleotide (ACO) via a space 9 linker, the compound was exposed to human serum for varying periods ranging from 24 hours to 14 days. The samples were then analyzed using polyacrylamide gel electrophoresis (PAGE) after trypsin digestion to strip bound proteins. As depicted in Supplementary Figure 1, from the first day onwards, RD-12500 progressively released its duplex siRNA, visible as a separate unbound duplex on the gel. In contrast, the ACO component mostly remained bound, indicated by the increased slow migrating bands on the top of the gel. This observation suggests that the siRNA-ACO conjugate maintained stability long enough to distribute within tissues, subsequently separating into two oligonucleotides and releasing the duplex for cellular functions. Additionally, this finding supports our hypothesis that the ACO component enhances the pharmacokinetics (PK) performance of the SCAD architecture.

#### Specificity and secondary pharmacology of RD-12500

Potential off-targets of RD-12500 was analyzed by *in silico* prediction and subsequently RT-qPCR evaluation of the expression of the predicted off-target genes in T98G cells treated with RD-12500 for 24 h. mRNA targets with perfect complementarity to the “seed” region of the RD-12500 guide strand (GS) and no more than 4 mismatches in the extended complementary region were selected as candidates. A total of 5 genes were identified including 4 with sites in 3’UTRs possessing 3-4 mismatches (*i.e*., CYTH2, NDUFC2, DISP2, and KNOP1) and one (*i.e.,* UBAP2L) with 2 mismatches in its coding region (Supplementary Table 1). RD-12500 at concentrations ranging from 0.25 nM to 25 nM had no significant effect on the mRNA level of CYTH2, NDUFC2, DISP2 and KNOP1 genes (Supplementary Figure 2A). For UBAP2L, a more detailed dose response assay revealed its expression was not significantly altered by RD-12500 at all concentrations tested (Supplementary Figure 2B).

#### Immunostimulation in PBMCs of RD-12500

Immune stimulatory activity of RD-12500 was assessed by ELISA assay in fresh human PBMCs treated with RD-12500 for 24 h by free uptake or RNAiMAX transfection. The siRNA duplex (RD-12928) and the ACO (RD-12340) components of RD-12500 were also included in the assay. As shown in Supplementary Figure 3. three selected cytokines (INF-α, TNF-α and IL-1β) did not show induction by RD-12500 and its two nucleic acid components (*i.e.,* the duplex and ACO).

### Biodistribution and clearance of IT administrated RD-12500 in rodents

To evaluate the distribution of RD-12500 in the CNS, we used RNAscope in situ hybridization (ISH) to detect the antisense strand of the siRNA in rat CNS tissues 14 days after a single intrathecal dose. The specificity of ISH assay was validated by using a scramble control probe in the brain section of RD-12500-treated rats and a RD-12500- specific probe in a blank brain section of the vehicle group (Supplementary Figure 4). The results (Figure 1) showed RD-12500’s extensive presence throughout the CNS, particularly in the spinal cord, especially its lumbar segment (Figure 1A), which is nearest to the injection site. Within the brain regions, the olfactory area, hippocampus, and cerebellum displayed higher levels of RD-12500 compared to other parts (Figure 1B).

**Figure 1.**
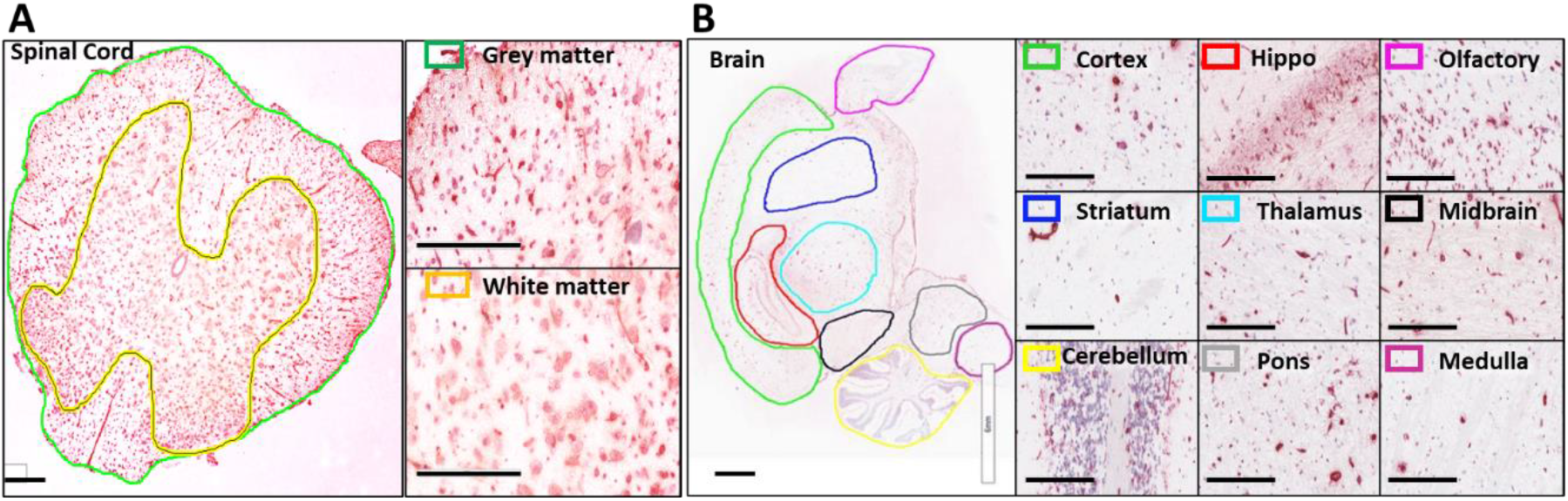
Tissue distribution of RD-12500 after single IT administration in rats. SD rats were dosed with 4.8 mg/dose of RD-12500 via IT injection and sacrificed on day 15. Harvested tissues were formalin-fixed and paraffin-embedded before subjecting to ISH. Representative images of RD-12500 in lumbar spinal cord (A) and the brain and brain regions (cortex, hippocampus, olfactory, stratum, thalamus, midbrain, cerebellum, pons and medulla) (B). Scale bar: 6 mm (whole brain) and 200 μm (spinal cord and in magnified regions). RNA signal is shown as red and cell nuclei were counterstained with hematoxylin.

We then analyzed the pharmacokinetic (PK) profile of RD-12500 in specific CNS and peripheral tissues of rats using liquid chromatography tandem mass spectrometry (LC-MS/MS). This study spanned from 6 hours to 42 days post a single intrathecal dose of 1.0 mg. The findings (Figure 2A and B) indicated that in the CNS, the spinal cord, especially the lumbar section, had higher drug levels than the brain regions, aligning with the biodistribution results (Figure 1 and Supplementary Table 2). Among the brain areas, the brain stem, cerebellum, hippocampus, and cerebrum showed comparable drug levels, while the striatum had the least. Notably, there was a rapid decline in drug concentration by day 7 post-dosing, which then stabilized for the remainder of the 42- day observation period (Figure 2C-F). Female rats exhibited slightly quicker CNS drug clearance compared to males (Figure 2C-F).

**Figure 2.**
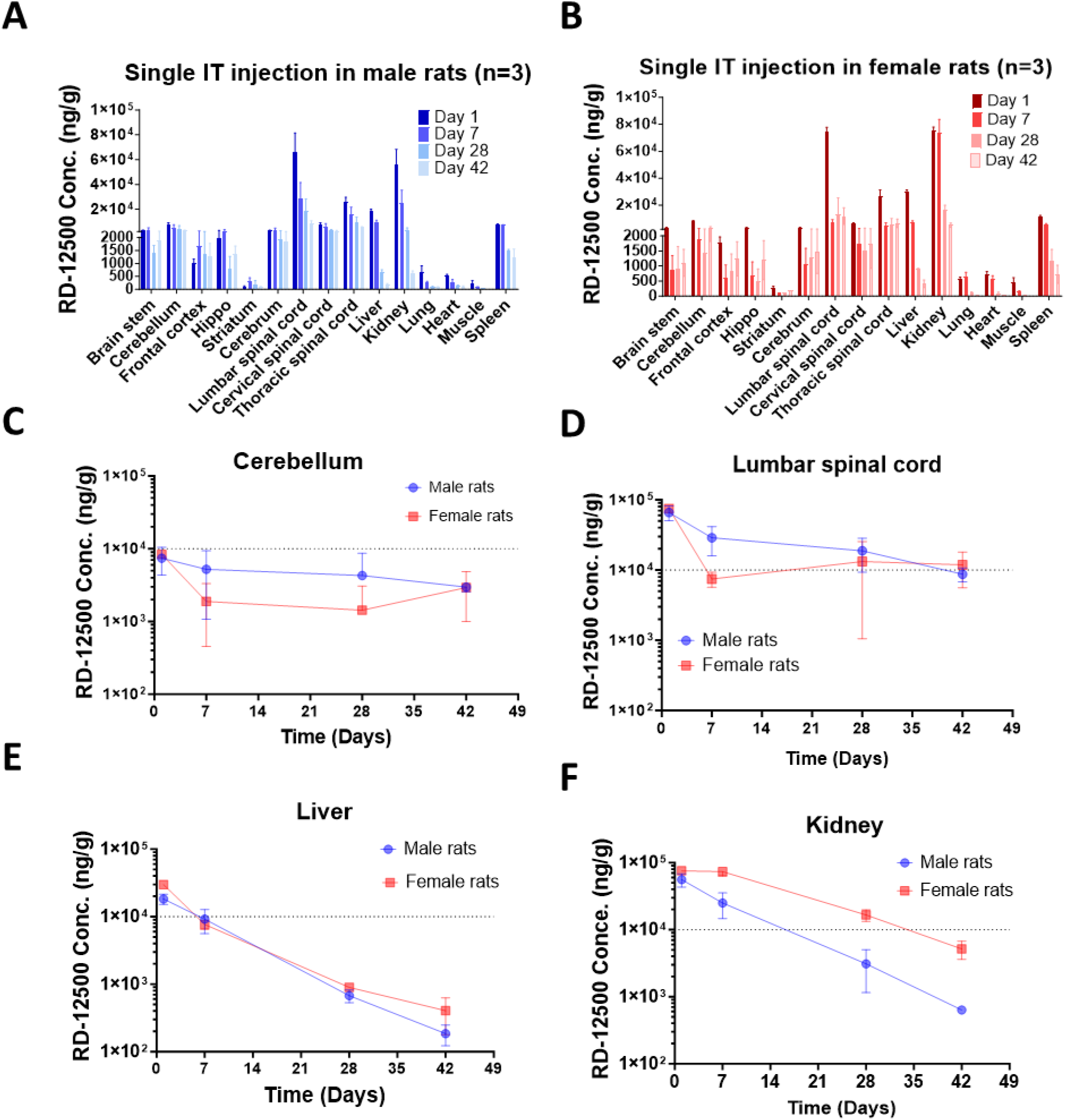
Tissue PK profile of RD-12500 in rats after single IT administration. Rats dosed with a single IT dose (1.0 mg/dose) of RD-12500 were sacrificed at different time points for tissue harvesting and subsequent siRNA concentration detection by LC-MS/MS. **A** and **B.** siRNA concentration in different tissues and at different timepoints in male (A) and female (B) rats. **C**-**F.** Representative PK profile in the brain (cerebellum, C), spinal cord (lumbar, D) and periphery tissues (liver and kidney, E and F). The dotted line in C-F denotes a siRNA concentration at 10000 ng/g.

Among the peripheral tissues studied, including the liver, kidney, spleen, lung, heart, and skeletal muscle, the liver, kidney and spleen showed exposure levels comparable to CNS tissues, with the kidney having the highest exposure (Figure 2A and B). However, peripheral tissues, notably the kidney and liver, demonstrated quicker drug clearance (half-life of 6-9 days) compared to CNS tissues (half-life of 20-27 days) (Figure 2C-F, Supplementary Table 2).

### Dose-and exposure-dependent deep and long-lasting knockdown activity of RD- 12500 in the CNS of cynomolgus monkeys

Sequence alignment of RD-12500 GS with cynomolgus monkey SOD1 mRNA revealed a mismatch at the 13th nucleotide from its 5’ end. Despite this, RD-12500 showed effective SOD1 knockdown in COS-1 cells (derived from an African green monkey with the same 13th nucleotide mismatch as cynomolgus monkeys), similar to its activity in T98G cells (Supplementary Figure 5). Consequently, cynomolgus monkeys were chosen for nonclinical toxicity and toxicokinetic studies.

Post IT administration at doses of 5, 20, and 50 mg, RD-12500 displayed dose-dependent exposure in the CSF and plasma of the monkeys (Supplementary Figure 6), with notably higher levels in the CSF. The CSF-to-plasma concentration ratios of RD- 12500 varied from 200-900 in the first 2 to 48 hours. The peak concentration in CSF was observed within 15 minutes post-dosing, followed by a rapid decrease within 48 hours, while plasma concentrations peaked between 2-4 hours and then decreased at a slower rate than in the CSF, indicating a drainage of the drug from CSF to plasma.

Pharmacodynamics analysis of tissue samples from the monkey study indicated that RD-12500 induced a dose-dependent reduction of SOD1 mRNA in CNS tissues (Figure 3B-E) and a decrease in protein levels in CSF (Figure 3F and G). At a 50 mg dose, significant SOD1 mRNA reduction (91%) was observed in the lumbar spinal cord and remained substantial up to 72 days. Moreover, a gradient decrease in RD-12500’s activity was noted, with the brain regions, being the furthest from the injection site, showing the least activity.

**Figure 3.**
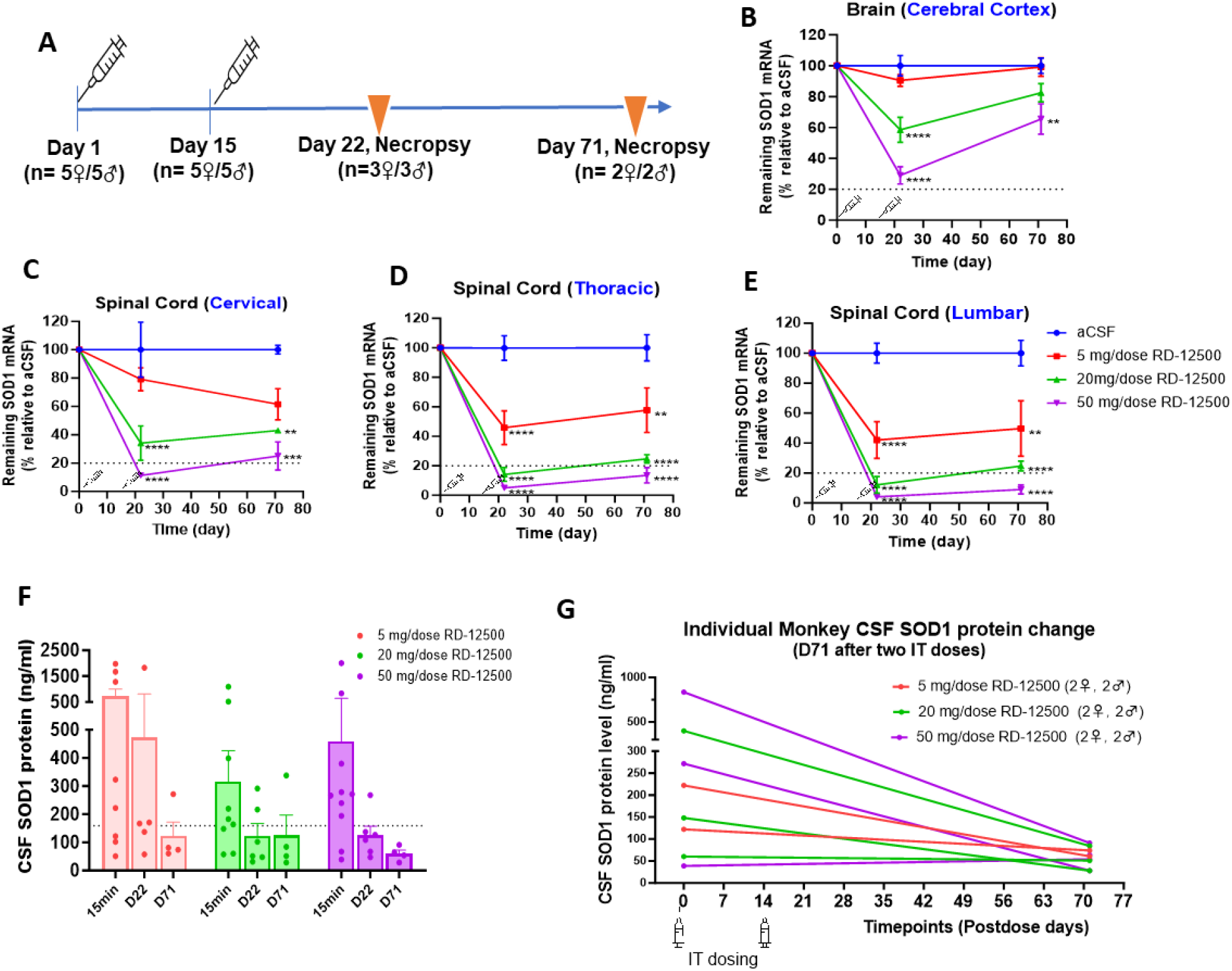
RD-12500 dose-dependently and durably lowered SOD1 mRNA expression in cynomolgus monkeys. **A.** Monkeys were dosed twice with aCSF or RD-12500 (5, 20, or 50 mg) via IT injection on day 1 and day 15, then necropsied for histopathology and incorporated pharmacology analysis at the indicated timepoints. **B-E.** SOD1 mRNA levels were quantified by RT-qPCR and plotted as the percentage of remaining SOD1 mRNA compared to aCSF group. Data represents mean ± SD on day 22 (n=6 per group) and on day 71 (n= 4 per group). **F** and **G.** SOD1 protein was assessed by ELISA assay and plotted as means ± SD (n = 6) of each group at different timepoints (F) and as individual data comparing day 71 and baseline (G). Statistical significance was determined by one-way ANOVA and Dunnett’s multiple comparisons test. *, *P* & 0.05; **, *P* & 0.01; ***, *P* & 0.001; ****, *P* & 0.0001.

### Pharmacokinetic/Pharmacodynamic relationship of RD-12500 in the CNS of cynomolgus monkey

An integrated analysis of both PD and PK data from cynomolgus monkeys indicated a robust dose-dependent effect of RD-12500. The estimated effective dose 50% (ED_50_) values for the cervical, thoracic, and lumbar spinal cord were 11.49 mg, 4.49 mg, and 3.86 mg, respectively (Figure 4A), and for the cerebral cortex, it was 50.23 mg (Figure 4D). The analysis also showed that SOD1 knockdown activity correlated with RD- 12500 tissue exposure (Figure 4B and 4E). The concentrations of RD-12500 predicted to achieve a median effective concentration (EC_50_) in the lumbar spinal cord and cerebral cortex were 158.2 ng/g and 374.5 ng/g, respectively, as determined by a 3- parameter logistic sigmoidal model (Figure 4C and 4F).

**Figure 4.**
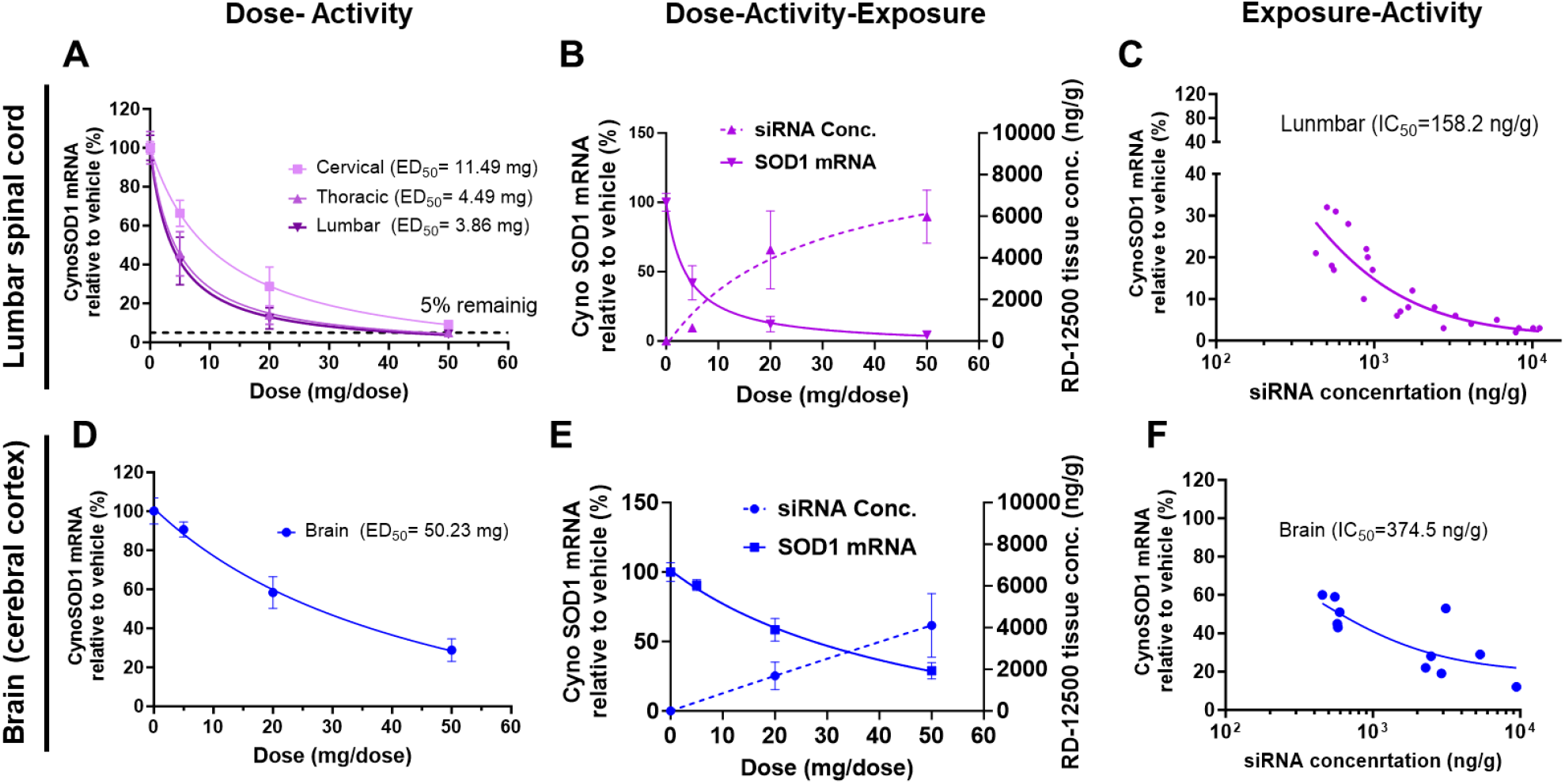
Pharmacokinetic and pharmacodynamic relationship of RD-12500 in the CNS of cynomolgus monkeys. **A** and **D.** Dose-dependent SOD1 knockdown in the spinal cord (A) and brain (D). **B** and **E.** Dose, exposure and activity relationship in the lumbar spinal cord (n=33) and frontal cortex (n = 21). mRNA knockdown activity and tissue concentration of RD-12500 are plotted on the left and right y axis respectively. **C** and **F.** Exposure and activity relationship in the lumbar spinal cord and frontal cortex.

### RD-12500 delayed disease onset, extended survival and improved mortal function in SOD1^G93A^ ALS rats

To verify RD-12500’s therapeutic efficacy, a repeat-IT dose study was conducted on SOD1^G93A^ ALS rats. Thirty-six of these rats were divided into two groups: one (n=16) received a scrambled duplex control (RD-13121), and the other (n=20) received RD- 12500. Both RD-13121 and RD-12500 had identical ACO conjugation and chemical modifications. The rats were given two IT doses of 0.9 mg on PND 70 and PND 130, with wildtype rats receiving two doses of aCSF as additional controls. The rats were monitored for changes in body weight, muscle strength, rotarod performance, neurological score, and survival. Typically, hemizygous SOD1^G93A^ rats show motor neuron disease onset between 174-207 days and reach end-stage disease about 11 days post symptom onset (12). Disease onset was defined by a neurological score of 1, marked by initial symptoms like abnormal hindlimb splay.

Figure 5 shows that the median disease onset in the RD-13121 group was 187.5 days, while the RD-12500 group had no median onset up to the study’s endpoint at 290 days. The median lifespan for SOD1 rats in the RD-13121 group was 215.5 days. Only 1 out of 16 RD-12500 treated rats showed disease onset, beginning at 205 days and reaching end-stage at 233 days by the study’s endpoint.

**Figure 5.**
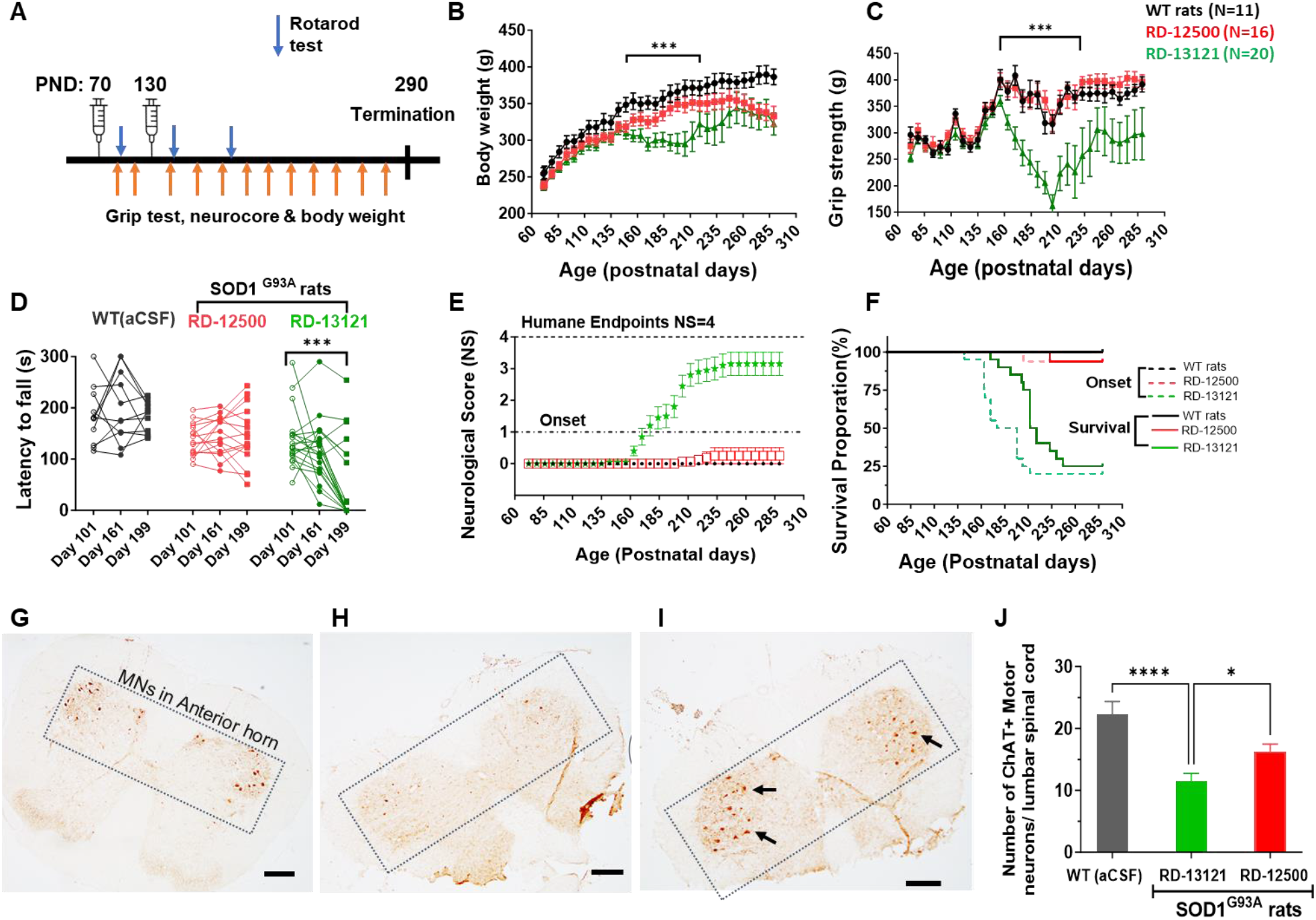
Therapeutic efficacy of RD-12500 in SOD1^G93A^ ALS model rats. **A.** Schematic of the study design. **B.** Body weight changes (%) over baseline. **C.** Muscle (hindlimb) grip strength. **D.** Neuroscore. **E.** Motor function as measured by rotarod test on PND101 (30 days after the first dose on PND70), PND 161 (30 days post-dosing on PND 70 and 130) and PND 199 (60 days post-dosing on PND 70 and 130). **F.** Log-rank (Mantel-Cox) analysis of median disease onset (NS ≥1) and lifespan (NS = 4, humane endpoint). **G-I**. MNs were stained with ChAT and counted within the hatched region of the cross-sectional L4-L5 segment in the lumbar spinal cord. Representative images of immunostaining with ChAT for the anterior horn are shown in G (Animal #16), H (Animal #45) and I (Animal #3). Arrows indicate ChAT-positive motor neurons. Bar is 100 μm. **J.** The total number of ChAT-positive neurons was counted and summed for each whole spinal cord. Data is expressed as mean ± SD of each group (WT: n = 3, RD-13121: n = 5 and RD-12500: n = 8).

The RD-13121 treated rats displayed disease onset around 160 days, marked by gradual weight loss, muscle weakening, deteriorating motor functions, and progressing paralysis. In contrast, RD-12500 treated rats maintained their body weight comparably to wild-type animals (Figure 5B) and showed improvements in muscle strength (Figure 5C), motor functions (Figure 5D), and delayed hindlimb paralysis (Figure 5E).

To investigate the underlying biological mechanisms of motor function improvement, motor neurons (MNs) in the lumbar spinal cord were identified using choline acetyltransferase (ChAT) staining and counted. Compared to wild-type rats, RD-13121 treated rats had a significantly reduced number of MNs, while RD-12500 treated animals had 42.2% more MNs than those treated with RD-13121 (Figure 5J).

### Late treatment of SOD1^G93A^ ALS mice with ICV-administrated RD-12500 significantly improved muscle strength and extended survival

Building upon our prior preclinical findings, where SOD1^G93A^ mice showed marked therapeutic response to early ICV (intracerebroventricular) and IT (intrathecal) administrations of RD-12500 (single dose on PND 85 or two doses on PND 68 and 100), we investigated the efficacy of delayed treatment, aligning with the typical post-symptomatic clinical intervention in ALS. SOD1^G93A^ mice were administrated with two ICV doses of RD-12500 on PND 126 and 151. For comparison, an early treatment group was included, receiving ICV doses at PND 70 and 100.

Aligning with our previous findings in SOD1^G93A^ mice (11), early treatment with RD- 12500 significantly enhanced motor function and muscle strength, stabilized body weight, and prolonged median lifespan to 331 days, a remarkable increase of 161.5 days (95%) compared to the vehicle control group.

Strikingly, late administration of RD-12500 also yielded significant benefits. Both low and high doses effectively enhanced motor function (Figure 6B), improved muscle strength (Figure 6C) and mitigated body weight loss (Figure 6D) in a dose-dependent manner. RD-12500 at 400 µg administrated at presymptomatic and presymptomatic delayed disease onset by 175 days and 45.5 days, respectively (Figure 6E). Compared to the aCSF control group, which had a median survival of 169.5 days, late-administered RD-12500 at both low (200 µg) and high (400 µg) doses significantly extended median survival by 78.5 and 128.5 days, respectively. This represents a significant increase in median survival of 46.3% and 75.8% for each dose (Figure 6F).

**Figure 6.**
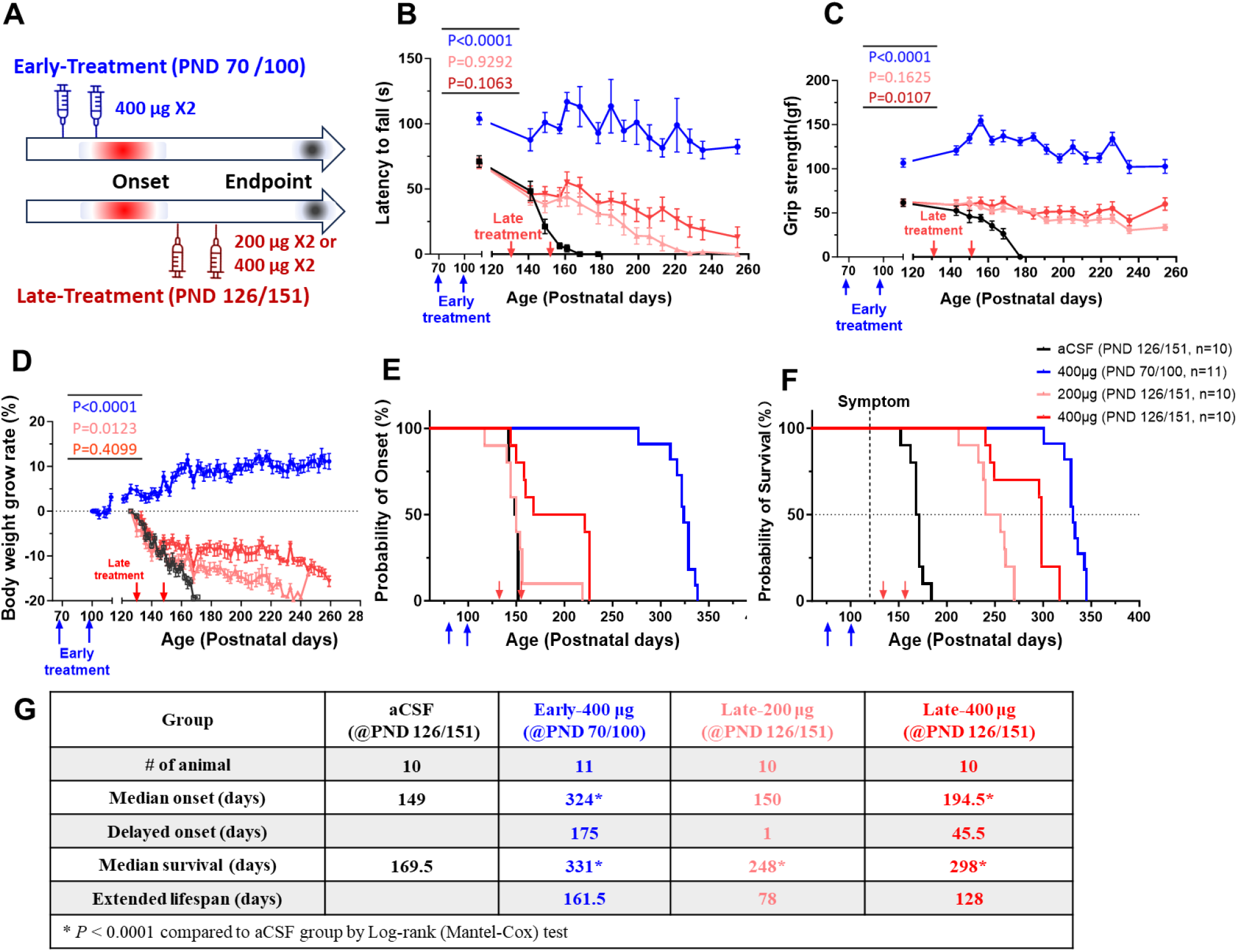
Therapeutic efficacy of early- and late-administrated RD-12500 in SOD1^G93A^ mice. SOD1^G93A^ mice were assigned into 4 groups: aCSF, early treatment group (Early-400µg) receiving 2 ICV doses of 400 µg RD-12500 at PD 70 and 100 days, late treatment low dose group (Late-200µg) and high dose group (Late-400µg) receiving 2 ICV doses of 200 µg and 400 µg, respectively, RD-12500 at PND 126 and 151 days. **A.** A scheme of study design of early or late-treatment in SOD1G93A mice. **B.** Mortal function as assessed by rotarod test. **C.** Muscle grip strength. **D.** Body weight change (%). **E.** Disease onset determined by 10% body weight loss relative to peak weight. **F.** Animal survival. Statistical significance was determined by one-way ANOVA and Dunnett’s multiple comparisons test compared with aCSF control. **G**. Summary of disease onset and survival data values present in (E) and (F). In A-C, *p* value in blue represents early-treatment, light and dark red represents respectively 200 μg and 400 μg late-treatment with RD-12500.

## DISCUSSION

Overcoming the challenge of effectively delivering duplex RNAs to the CNS has proven elusive, despite the myriad druggable targets accessible via duplex RNAs for treating various disorders, including neurodegenerative diseases. In this study, we employed an innovative delivery technology known as SCAD, comprising a duplex and an accessory oligonucleotide (ACO) linked together. This approach facilitated the local injection delivery of a siRNA-ACO conjugate, RD-12500, targeting SOD1, resulting in extensive distribution across CNS tissues and achieving sustained and substantial reduction in SOD1 levels. Notably, the administration of RD-12500 to SOD1^G93A^ ALS mice and rats demonstrated remarkable therapeutic efficacy, even when treatment was initiated post disease onset.

SOD1^G93A^ model is one of the most widely published transgenic mouse models for ALS. Typically, researchers start treatment early to better magnify and elucidate etiological impact (13). In a survey study (13), Abati et al. analyzed more than 4000 SOD1^G93A^ transgenic ALS mouse data points and found that the mean treatment start date for the presented analysis was postnatal 49.8 ± 1.5 days, which was well before symptom onset in SOD1^G93A^ mice and less than 5% of ALS preclinical therapies started after functional symptom onset. The ENMC (European Neuromuscular Centre) recommends to conduct proof of concept studies between day 50 and day 70 (in the G93A B6SJL animal model) given the high copy number expressing and highly aggressive nature in this model (14, 15). In a review article (16), Carrì et al summarized 6 publications in which treatment was given to SOD1^G93A^ mice at or after onset of symptoms. In the 6 studies, no treatment was given after PND 90, and the highest increase in lifespan was 22% (by 24 days) (17) or by 17 days (11%)(18).

In our recent published studies, we have demonstrated the remarkable therapeutic efficacy of RD-12500, when administered before disease onset (PND 68 and PND 85), in SOD1^G93A^ mice (11), a model on pure C57BL/6J congenic background (Stock No. 004435) for ALS with typical onset between 90 to 100 days and endpoint from 147 to 166 days (15, 19). Based on such encouraging data, we wished to explore potential therapeutic benefits in the same animals by treating them after disease onset. We dosed SOD1^G93A^ mice twice on PND 126 and 151 (late treatment group) when animals had already reached endpoint. Despite such delayed treatment, the late treated animals performed significantly better than aCSF treated mice across all metrics evaluated including body weight maintenance, motor function and muscle strength improvement, and extension of survival. This preclinical efficacy date may support potential therapeutic benefit of RD-12500 not only for pre-symptomatic SOD1-ALS patients but also post-symptomatic or even patients in late stages of the disease. To the best of our knowledge, this is the first study to demonstrate the therapeutic benefits of a treatment administered well after the onset of ALS symptoms in SOD1 ^G93A^ ALS mice.

The serum stability analysis of RD-12500 provides crucial insights into its pharmacokinetic behavior. The study’s results, showing a gradual release of the siRNA duplex from the ACO component, highlight the stability and controlled release properties of RD-12500 in biological environments. This suggests that the drug can remain intact long enough to be distributed within target tissues before releasing its active components, a characteristic important for ensuring therapeutic efficacy.

The PK/PD relationship established in the study will be hopeful in guiding clinical dosing strategies. The dose-dependent effect of RD-12500 and the correlation between drug exposure and SOD1 knockdown provide a quantitative basis for determining effective dosing levels in patients while minimizing risks. The sustained SOD1 knockdown effect of RD-12500 over two months, as observed in monkey studies, is particularly noteworthy when compared to recently FDA-approved Tofersen which requires dosing once every 28 days (20). The long-lasting PD effect of RD-12500 suggests a potentially more extended interval between doses compared to Tofersen, which could offer significant advantages in terms of patient compliance, overall treatment management and safety. Such a pharmacological profile is especially beneficial in managing chronic conditions like ALS, where treatment adherence and minimizing patient burden are key considerations.

In conclusion, our findings demonstrate that even when administered at late disease stage, RD-12500 can effectively delay ALS progression, improve neuromuscular function, and extend lifespan. This positions RD-12500 not only as a viable alternative to existing ASO therapies but also highlights its particular efficacy in situations where treatment initiation is unavoidably delayed, addressing a significant challenge in ALS patient care. Additionally, an investigator-initiated trial (IIT) is currently underway, aiming to evaluate the safety and tolerability of RD-12500 (RAG-17) in ALS patients with the SOD1 gene mutation (ClinicalTrials.gov Identifier: NCT05903690), marking a crucial step in translating our findings into clinical practice.

## MATERIALS AND METHODS

### Oligonucleotides synthesis

Oligonucleotides was synthesized in house for in vitro studies and efficacy studies in mice and rats as previously described (21). Briefly, single strand oligonucleotide was synthesized on solid-phase support using a HJ-12 synthesizer (Highgene-Tech Automation, Beijing, China) and subsequently purified via RP-HPLC using an acetonitrile gradient over a UniPS column (NanoMicro Technology, Suzhou, China). Each sequence was reconstituted in sterile water via buffer exchange. Equal molar quantities of each strand were annealed to make a duplex by briefly heating the strand mixtures and then cooling to room temperature. Resolution of a single band via gel electrophoresis at predicted molecular weights was used to qualify duplex formation. ESI-MS was used to confirm duplex identity, while overall purity was analyzed via SEC-HPLC using a XBridge Protein BEH SEC 125 A column (Waters Corporation, Milford, MA, USA). Endotoxin levels in each batch were quantified using the end-point Chromogenic Endotoxin Quant Kit (Bioendo, Xiamen, China) via proenzyme Factor C. Oligonucleotide (RAG-17) used for NHP toxicology study synthesized by Wuxi SynTheAll Pharmaceutical Co., Ltd (Lot# TT-22E029).

### Serum stability assay

siRNA or siRNA-ACO duplexes were incubated at 37°C with PBS or 50% active human serum at a final concentration of 3 μM for different durations from day1 to day14. The oligonucleotides-proteins complexes were then treated with 20 mg/ml trypsin for 30 min at 37°C. Ten microliter of each sample was loaded to a 20% TBE-PAGE-UREA gel and electrophoresed at 120 V for 90∼120 min. Gel images were captured on the ChemiDoc XRS+ Imager System (Biorad, Hercules, CA, USA).

### *In silico* off-target prediction

*In silico* off-target prediction of siRNA sequences were performed using the *in silico* alignment algorithm from gggenome (https://gggenome.dbcls.jp) and validated with in silico search algorithm miRanda (22) against all human mRNA sequences retrieved from the Ensembl genome database. The output validation results were parsed using an in-house script which also tabulated the output based on target site location in mRNAs (i.e., 3’UTR or coding region). Candidates found with GGGenome and validated with miRanda, or found uniquely with miRanda were furthered scored based on the number of matches and mismatches in each region (*i.e*., seed or flanking region). Wobbles, a form of non-standard Watson-Crick base pairings between G and U nucleotides were classified as matches. Exemplary off-target candidates with overall high complementarity including 4 with sites in 3’UTRs possessing 3-4 mismatches (i.e., CYTH2, NDUFC2, DISP2, and KNOP1) and a single transcript (i.e., UBAP2L) with 1-2 mismatches in its CDR were selected from the *in silico* results to experimentally evaluate RD-12500 specificity by RT-PCR assay.

### Cell culture and transfection

293A cells (Cobioer, Nanjing, China, Cat# CBP60436) and SK-N-AS (Procell, Wuhan, China, Cat# CL-0621) cells were maintained in DMEM medium supplemented with 10% FBS, penicillin (100 U/ml) and streptomycin (100 mg/ml). T98G cells (Cobioer, Cat# CBP60301) were maintained in MEM medium supplemented with 10% FBS, 1% NEAA, sodium pyruvate (1 mM), penicillin (100 U/ml) and streptomycin (100 μg/ml). All cell lines were cultured in a humidified atmosphere of 5% CO_2_ at 37°C. Transfections were carried out using Lipofectamine RNAiMax (ThermoFisher, Waltham, MA, USA) in growth media without antibiotics according to the manufacture’s protocol.

### *In vitro* immunotoxicity assay in PBMC cells

Heparinized fresh blood from three different health volunteers was diluted with an equal volume of PBS. Peripheral blood mononuclear cells (PBMC) were separated by density gradient centrifugation, using Ficoll-Paque (Cytiva 17144003) in Leucosep filter tubes (Serymwerk 11421) at 2000 rmp for 10 min at room temperature. The interface layer consisting of mononuclear cells was collected and washed once with PBS and once again with cell culture media. PBMC were resuspended in culture medium consisting of RPMI 1640 medium and 1 million /well PBMC was seed in 96 round well plate for overnight. For free uptake, nontreatment replicate wells served as a blank control, 1 μg /mg lipopolysaccharides (LPS) and 1 μM CpG ODN2216 was incubated for 24-h as positive controls. Two replicate wells were exposed to two concentrations (1 μM or 10 μM) of each of the 3 test articles. For RNAiMAX transfection, RNAiMAX alone was used for mock control, 133 nM RAG-IS-1 and CpG OND2216 were used as positive controls, each test article was transfected at 33 or 133 nM final concentration with 1 μl RNAiMAX for 24 h. The supernatants were for determination of cytokine release. Cytokine levels in the supernatants were quantified using Human IFN-α ELISA kit (Cat. # EK182-96, Lot.: A18220621), Human TNF-α ELISA kit (Cat. #EK182-96, Lot.: A18220621), Human TNF-α ELISA kit (Cat. #EK182-96, Lot.: A18220621) according to the protocol of the manufacturers.

### Gene expression analysis via RT-qPCR

Animal tissue frozen in RNALater (Sigma-Aldrich, St. Louis, MO, USA) was homogenized in Total RNA Isolation Reagent (Biosharp, Hefei, China) using a Bioprep-24 Homogenizer (Allsheng, Hangzhou, China). Chloroform was added to the homogenate in which the aqueous phase was removed and mixed with isopropanol. Total RNA was extracted from the tissue prep using the RNeasy RNA kit (Qiagen) according to the manufacture’s protocol. RNA from cell culture was extracted using the Auto-Pure 96A (Allsheng) nucleic acid extraction system. Reverse transcription (RT) reactions were performed with 1 μg total RNA using the PrimeScript RT kit with gDNA Eraser (Takara). The resulting cDNA was amplified in triplicate on the 480 Real-Time PCR system (Roche) using SYBR Premix Ex Taq II (Takara) in conjunction with primer sets specific to mRNA of test genes and an internal control for either human (*i.e.*, TBP) or mouse (*i.e.*, mTbp) samples. Melting curves were performed after amplification to confirm primer specificity. Averaged Ct values for each sample were used to calculate relative gene expression via ΔΔCt method. All PCR primer sequences are listed in **Supplementary Table 3**.

### Animal handling and grouping

Parental transgenic hSOD1^G93A^ mice (Strain ID #004435) were purchased from The Jackson Laboratory (Bar Harbor, ME, USA) and imported into China via Nantong University (Nantong City, Jiangsu Province, China). Mice were delivered to the animal facility at 6 weeks of age and subsequently bred domestically at Nantong University who supplied the animals for this study. All animal procedures were approved by the IACUC at Nantong University. Formulations for animal treatments were prepared fresh prior to use by dissolving allotments of lyophilized oligonucleotide into aCSF to create stock solutions for dilution to the intended treatment concentrations. Animals were randomly allocated into study groups based on body weight. Any animals in poor health or with obvious abnormalities were omitted from the experiments. Randomization was analyzed using Ordinary one-way ANOVA via GraphPad Prism version 8.3.0 Windows (GraphPad Software, San Diego, CA, USA).

### Intracerebroventricular (ICV) injection

Avertin (1.2%) was prepared fresh and sterilized via 0.2-micron filter. Mice were dosed at 0.30-0.35 ml per 10 g body weight via intraperitoneal (IP) injection in a stereotaxic apparatus to rapidly induce anesthesia for up to 30 minutes. An approximate 11.5 mm incision was made in the animal’s scalp and a 25-gauge needle attached to a Hamilton syringe containing the appropriate siRNA formulation was placed at bregma level. The needle was moved to the appropriate anterior/posterior and medial/lateral coordinates (0.2 mm anterior/posterior and 1 mm to the right medial/lateral). A total of 10 μL was injected into the lateral ventricle at an approximate rate of 1 μL/s. Following treatment, the needle was slowly withdrawn, and the wound sutured close.

### Intrathecal (IT) injection

Rats were anesthetized via 3.0% isoflurane in an induction chamber for a continuous 10 mins. Hair was shaved around the injection site at the base of the tail and cleaned with 75% ethanol. The space between the L5-L6 spinous processes was identified and a 30-gauge needle attached to a microliter syringe containing the appropriate drug formulations was slowly inserted into the intradural space until a tail flick was observed. The needle position was subsequently secured in which 30 μL total volume of solution was injected over the course of 1 min.

### Efficacy study in SOD1^G93A^ transgenic rats via IT administration

SOD1^G93A^ transgenic rats (23) (Taconic Biosciences, model 2148, provided by Cyagen) were anesthetized via 3.0% isoflurane in an induction chamber for a continuous 10 mins and then received a 30-μl intrathecal bolus injection between the L5 and L6 vertebrae of the lumbar spinal cord. Hair was shaved around the injection site at the base of the tail and cleaned with 75% ethanol. SOD1 rats aged of postnatal 70 days were given 1^st^ dosing of 900 μg RD-13121 or 900 μg RD-12500 (the date to 1^st^ dosing was defined Day 0) and 2^nd^ dosing Day 60 (postnatal 130 days). WT rats from the same litter were used for health control and received two IT doses with an artificial cerebrospinal fluid (aCSF) on Day 0 and Day 60. In addition, technicians were blinded to the identity of the RD-13121 and RD-12500 at the time of treatment and throughout the duration of the experiment. The study protocol was approved by Institutional Animal Care and Use Committee (WuXi Apptec, IACUC #: GP02-QD105-2020V1.0)

### Efficacy study in SOD1^G93A^ transgenic mice via ICV injection

Parental transgenic SOD1^G93A^ mice (B6.Cg-Tg(SOD1*G93A)1Gur/J, Strain ID #004435) were purchased from The Jackson Laboratory (Bar Harbor, ME, USA) and imported into China via Nantong University (Nantong City, Jiangsu Province, China). Mice were delivered to the animal facility at 6 weeks of age and subsequently bred domestically at Nantong University who supplied the animals for this study. All animal procedures were approved by the IACUC at Nantong University (#S20220323- 001). The animals were anesthetized via fresh prepared and filtered 1.2% avertin and then received 10 uL/dose lateral intracerebroventricular (ICV) injection (0.2 mm anterior/posterior and 1 mm to the right medial/lateral away from the bregma) at an approximate rate of 1 μl/s. For early-treatment regimen, SOD1^G93A^ transgenic mice were dosed with vehicle (aCSF) or 400 μg of RD-12500 on postnatal 70-days and 100- days. For late-treatment regimen, 200 or 400 μg of RD-12500 was dosed in two additional groups of SOD1^G93A^ transgenic mice on postnatal 126-days and 151-days.

### Clinical observation and endpoint criteria

Cage-side clinical signs, *i.e.,* ill health, behavioral change et.al. were recorded at least once daily during the acclimation period and twice a day (AM and PM) during the treatment. If abnormal symptoms occur, describe them in detail and collect videos or take photos. Such as convulsions, seizures, rigidity, muscle tremor, and dyspnea. Animals were observed after injection for up to 4 h and daily thereafter until endpoint. Body weight was recorded before test substance administration and at recorded intervals thereafter. Animals with weight loss >20% relative to their initial mass at day of treatment or a Neuroscore = 4 met humane endpoint criteria.

### Neurological scoring (Neuroscore)

For animals treated via IT injection, mice were evaluated for signs of motor deficit using the ALS Therapy Development Institute (ALS TDI) neuroscore (NS) system, which was developed to provide unbiased assessment of disease progression based on hindlimb dysfunction common to SOD1^G93A^ mice (24). NS was assigned based on the following 4-point scale: 0 if no signs of motor dysfunction (*i.e.*, pre-symptomatic), 1 if hindlimb tremors are evident when suspended by tail (*i.e.*, first symptoms), 2 if gait abnormalities are present (*i.e.*, onset of paresis), 3 if dragging at least 1 hindlimb (*i.e.*, partial paralysis), and 4 if inability to right itself within 10 seconds (*i.e.*, endpoint paralysis).

### Grip strength test

Mice were lowered onto a grid plate in which its forepaws and hind paws were allowed to grasp the grid. The tail was gently pulled and maximal muscle strength was measured on the XR501 grip strength meter (XinRuan Information Technology) in units of mass until the animal relinquished its grasp. Each animal was tested in triplicate in which the mean value represented grip strength. Grip strength assessment in rats was carried out using a digital grip-strength meter equipped with a bar (BIOSEB, Vitrolles, France, model BIO-GS3).

### Rotarod test

After pre-training of rats at rotating at 4 rpm in 5 min, Mice were placed on a motionless rotarod apparatus (Harvard Apparatus, Panlab Rotarod, LE8305, USA) with a swivel bar 60 mm in diameter. Rotational speed was accelerated from 0-40 rpm over the course of 300 seconds. Latency time was recorded as the amount of time it took for each animal to fall off the swivel bar. Each animal was tested in triplicate in which the longest value represented latency time.

### RNAscope *in situ* hybridization

SD rats were IT dosed with RD-12500 at 1.6 mg/dose or 4.8 mg/dose or saline as vehicle control. The brain and spinal cord were harvested and paraffin-embedded tissue sections were prepared for in situ RNA hybridization detection by RNAscope assay according to the manufacturer’s protocol (Advanced Cell Diagnostics) of the miRNAscope™ HD Reagent Kit - RED (ACDbio-324500, California, US) on paraffin slides. siRNA. The antisense strand of RD-12500 was detected by a customized probe (ACDbio-1199511-S1). A scramble control probe (ACDbio-727881-S1) and positive control RNU6 probe (ACDbio-727871-S1) were also used.

### Tissue distribution and quantification of RD-12500 in rats

Thirty-six (18/sex) SD rats were divided into six groups with 3 animals/sex/group and received a single intrathecal dose of RD-12500 at 1.0 mg/dose with a 30-μl intrathecal bolus injection between the L5 and L6 vertebrae of the lumbar spinal cord. Six (3/3) rats were euthanized by CO_2_ inhalation each of the following time points: 0.25 (CSF only), 2 (CSF only), 6 (Day 1), 144 (Day 7), 648 (Day 28) and 984 (Day 42) h post-dose. Brain tissues, spinal cord and periphery tissues were collected and homogenized. Each tissue for liquid chromatography tandem mass spectrometry (LC-MS/MS) analysis was homogenized with a 1:9 (w/v) ratio of homogenizing buffer on wet ice (1 g tissue with 9 mL of homogenizing buffer) using pre-cooled Lysis-loading buffer (phenomenex, clarity OTX). About 0.8 mL of tissue homogenate samples (if enough) were transferred to a low-binding tube. The samples were stored at -60°C or lower until analysis. The quantitative determination of RD-12500 in SD rat brain homogenate by a sensitive, specific and reproducible LC-MS/MS method which was validated in brain homogenate and liver homogenate. The dynamic range of the method was 50.00- 25000 ng/g

### Concentration of RD-12500 in plasma, CSF and tissues of cynomolgus monkey

Monkey Samples (Plasma, CSF and selected tissues) were derived from nonclinical safety study which was conducted in WuXi AppTec (Suzhou) Co., Ltd. (Study #: H59- 0027-TX). Briefly, 40 monkeys (20/sex) aged 3 to 4 years were assigned randomly to 4 groups of 5/sex/group and received 2 IT doses, on day 1 and day 15, of artificial CSF as vehicle control or RD-12500 at dose levels of 5, 20, or 50 mg/dose.

#### Plasma Samples

Blood samples (approximately 0.5 mL in K2EDTA) were collected at 0 (predose), 0.25 (15 minutes), 0.5, 2, 4, 6, 8, 12, 24, and 48 h postdose from all animals. Plasma was obtained within 1 hour of collection by centrifugation at 3200×g and 4°C for 10 minutes and stored in a freezer set to maintain < 60°C until transferred and analysis.

#### CSF samples

Cerebrospinal fluid (CSF, approximately 0.3 mL) was collected from all animals once during pretest and at 0.25 (15 minutes ± 3 minutes), 24, 48, 72, and 168 h postdose from lumbar vertebrae or cisterna magna. The volume ratio of CSF to stabilizer was 1:1. The tubes were gently inverted several times to ensure mixing and immediately placed upright on dry ice, and stored in a freezer set to maintain at <-60°C until transferred and analysis.

#### Tissue samples

3/sex/group and 2/sex/group monkeys were sacrificed on day 22 and on day 71. The brain, spinal cord and liver samples were collected. The tissue samples were weighted, washed with cold saline, snap frozen in liquid nitrogen for PK and PD analysis.

### Quantification of RD-12500’s concentration by LC-MS/MS

The quantification of RAG-17 concentration was conducted by using a validated liquid chromatographic triple quadrupole mass spectrometric (LC-MS/MS) method. The linearity for RAG-17 20.0 to 8000 ng/mL and 40.0 to 16,000 ng/mL in plasma and CSF, respectively. Monkey brain or spinal cord homogenate was prepared in a ratio of 1:9:10 between spinal cord tissue, de ionized water and monkey plasma. The preparation process was weight about 1.00 g spinal cord tissue, add 9.0 mL of de-ionized water and 10.0 mL monkey plasma, mix well. The lower limit of quantification (LLOQ) for RAG- 17 was 20.0 ng/mL (400 ng/g) and the upper limit of quantification (ULOQ) was 8,000 ng/mL (160,000 ng/g). Corrected Concentration=Concentration×20. All bioanalytical data were collected by Analyst™ and processed using Watson LIMS.

### Cynomolgus monkey Sod1 mRNA expression in tissue detected by RT-PCR

Frozen animal tissue in RNALater (Sigma-Aldrich) was homogenized in RNAVzol using a Bioprep-24 Homogenizer (Allsheng). Chloroform was added to the homogenate in which the aqueous phase was removed and mixed with isopropanol. Total RNA was extracted from the tissue prep using the RNeasy RNA kit (Qiagen) and reverse transcribed into cDNA using xx. The resulting cDNA was amplified by quantitative PCR. Each sample was amplified in triplicate on the 480 Real-Time PCR system (Roche) using SYBR Premix Ex Taq II (Takara) in conjunction with primer sets specific to cynoSOD1 and an internal control (i.e., cynoGAPDH) as **Table S2**. Melting curves were performed after the amplification cycles were complete to confirm primer specificity. Averaged Ct values for each sample were used to calculate relative gene expression via ΔΔCt method. Percent (%) knockdown was calculated as 1-(relative SOD1 levels) *100.

### SOD1 protein quantification in the CSF of monkeys

SOD1 protein levels in the CSF of monkeys were measured using Monkey Superoxide dismutase 1, soluble (SOD1) ELISA Kit (Cat# AE17754MK, Abebio) specifically qualified to quantify the amount of SOD1 in cynomolgus monkey CSF. Briefly, serial diluted standards or CSF samples were pipetted into pre-coated microplate and allowed to incubate at 37°C for 2 hrs. After removing any unbound substances, a biotin-conjugated antibody specific for SOD1 was added to the wells at 37°C for 1 hr. After washing, streptavidin conjugated horseradish peroxidase (HRP) was added to the wells at 37°C for 1 hr. Following a wash to remove any unbound avidin-enzyme reagent, a substrate solution was added to the wells and the plate was read in a microplate reader at absorbance 450 nm with the correction wavelength set at 540 nm.

### Statistical analysis

Data analytics were performed using GraphPad Prism version 8.3.0 Windows. The differences in continuous variables between treatments were assessed by two tailed and unpaired Student t test (for 2 treatments) or one-way ANOVA followed by Tukey’s multiple comparison test (for 3 or more treatments). Time-stratified data (*e.g.*, peak weight analysis and animal survival) was plotted via Kaplan-Meier graphs in which statistical significance was verified using the Mantel-Cox test. Significance was defined as *P* < 0.05.

