## Supplementary Figures and Tables for "RAG-17: A Novel siRNA Conjugate Demonstrating Efficacy in Late-Stage Treatment of SOD1^G93A^ ALS mice"

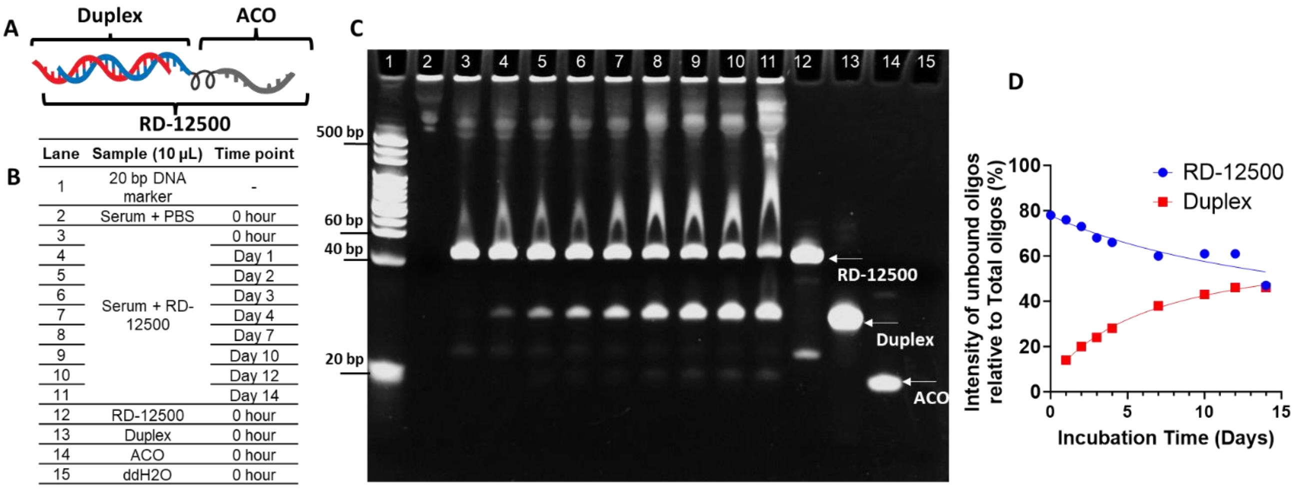


**Supplementary Figure 1. Stability of RD-12500 in human serum. A.** Schematic of SCAD architecture that includes a duplex conjugated to an ACO via a linker. **B** and **C**. Oligonucleotides (siRNA-ACO, duplex siRNA, ACO) at 3 µM were incubated with equal volume of active human serum (final concentration of 50% v/v) at 37°C for different duration and separated by PAGE. Sample mixed with PBS as 100% unbound control (lane 12-14). Lane information of samples loading is shown in (B). **D.** Relative band intensity in (C) for RD-12500 band and free siRNA duplex band.

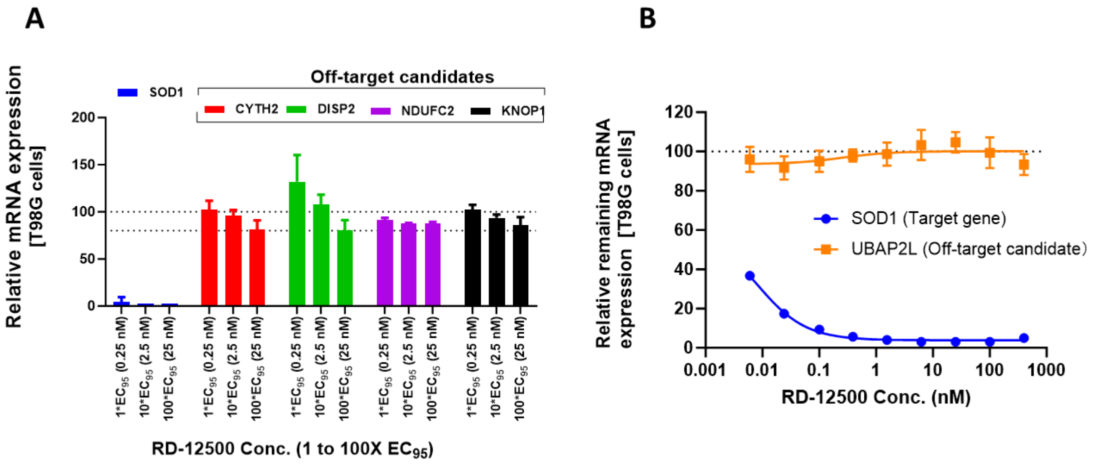


**Supplementary Figure 2. RT-qPCR assessment of potential off-target gene expression**. **A.** RT-PCR analysis following transfection with RD-12500 at concentrations up to 100x its IC95 for on-target activity (i.e., 0.25, 2.5, and 25 nM) of high complementarity possessing 3-4 mismatches in 3’UTRs (i.e., CYTH2, NDUFC2, DISP2, and KNOP1). **B**. A complete dose response curve for UBAP2L expression which carries 1-2 mismatches in its CDR.

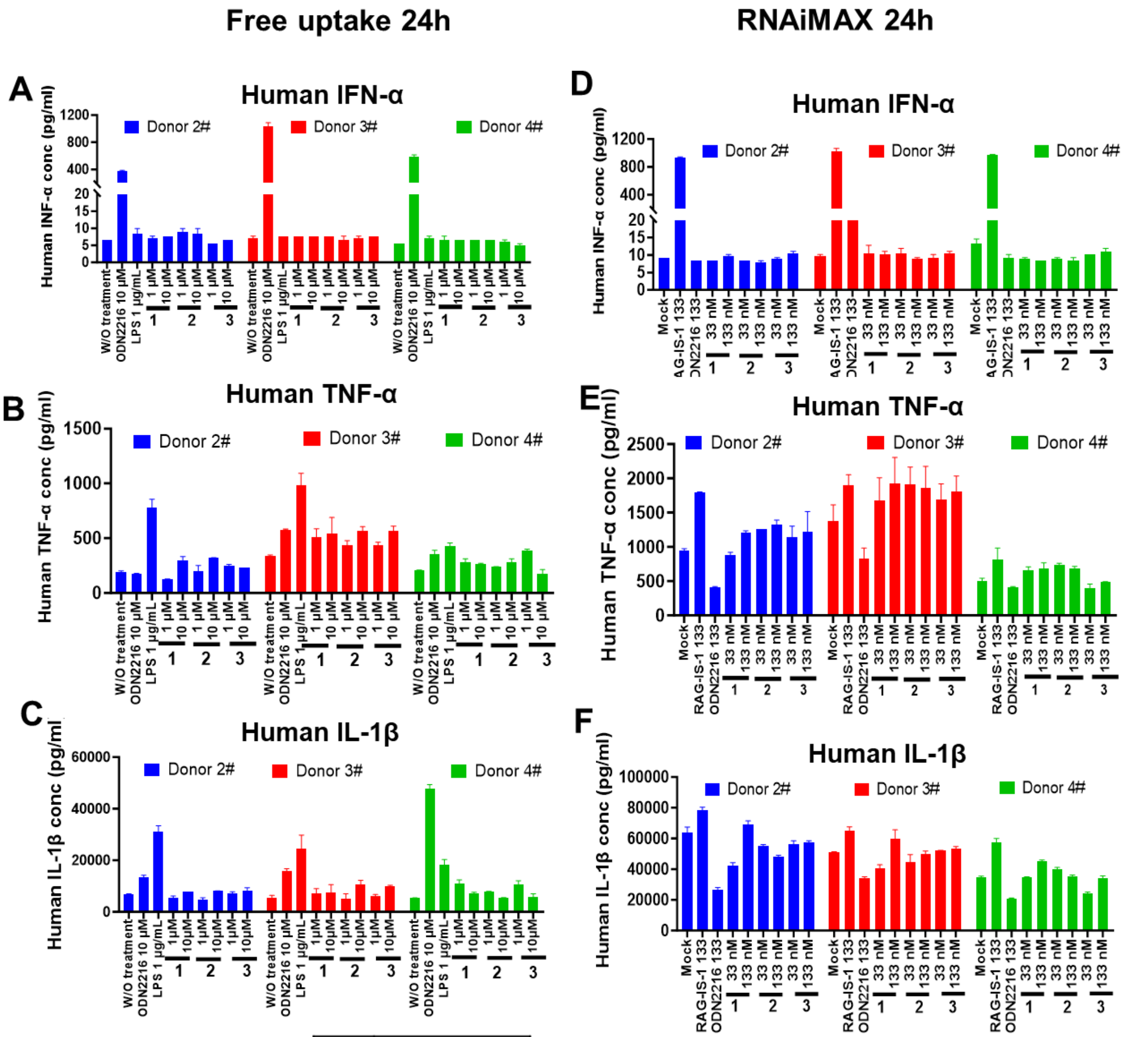


**G**

| | |
|---|---|
| 1 | RD-12500 |
| 2 | RD-12928 |
| 3 | RD-12340 |

**Supplementary Figure 3. PBMC-based immunotoxicity screen. A-C.** Oligonucleotides (as listed in G) were added to freshly isolated PBMC from 3 healthy donors at 1 μM or 10 μM and 24 h later, INF-α, TNF-α and IL-1β was measured from the supernatants by ELISA. **D-F**. Oligonucleotides (as listed in G) were transfected using RNAiMAX into freshly isolated PBMC from 3 healthy donors at 33 nM or 133 nM and 24 h later, INF-α, TNF-α and IL-1β was measured from the supernatants by ELISA. Mock was transfected in the absence of oligonucleotides. CpG ODN2216 (ODN2216) and LPS served as positive controls, nontreatment (W/O) served as a blank control, RD-12928 and RD-12340 are respectively the duplex and ACO components of RD-12500. **G**. The code name of oligonucleotides tested in the assay.

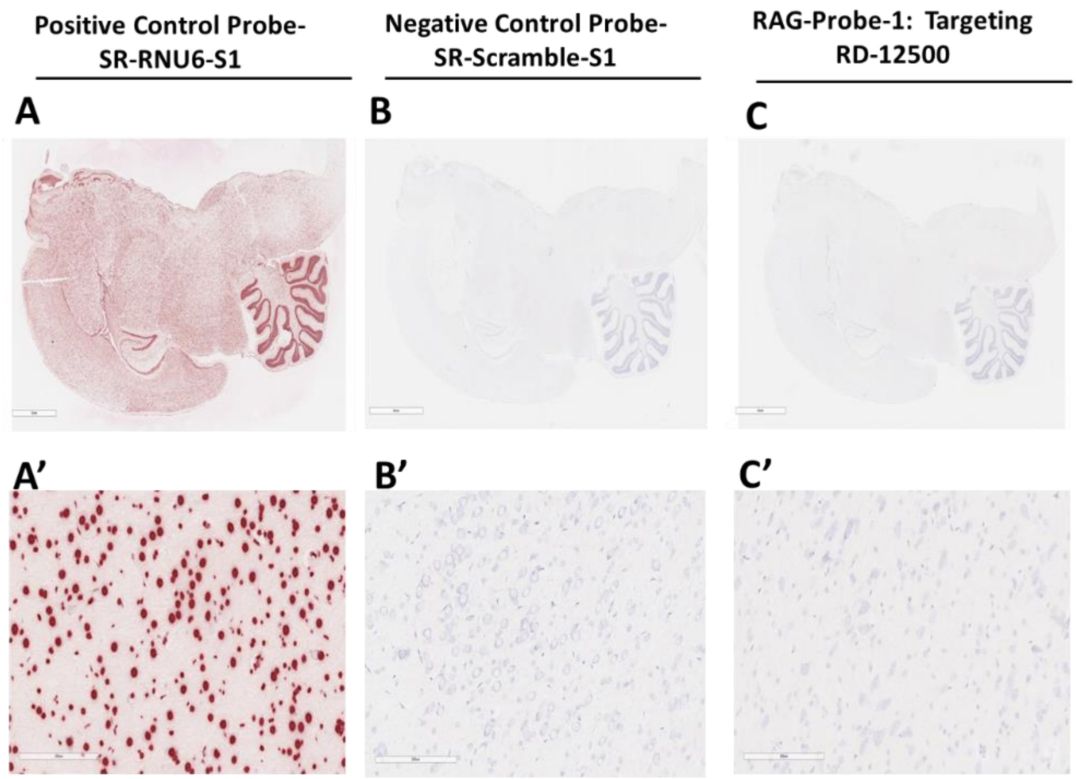


**Supplementary Figure 4. Specificity of RNAscope assay.** (A-B) Positive control probe [SR-RNU6-S1 (727871-S1] which hybridizes with U6 small nuclear 1 RNA) and negative control probe (SR-Scramble-S1, 727881-S1, a probe with scrambled sequence) were used to validate specificity and quality of assay in paraffin-embedded brain section of rats from vehicle group. (C) RD-12500 probe against blank brain sections from vehicle (aCSF) control group rats. A’-C’ is enlarged image of A-C, respectively. The scale bar is 6 mm for the whole brain and spinal cord and 200 µm in magnified regions. RNA signal is shown as red and cell nuclei were counterstained with hematoxylin.

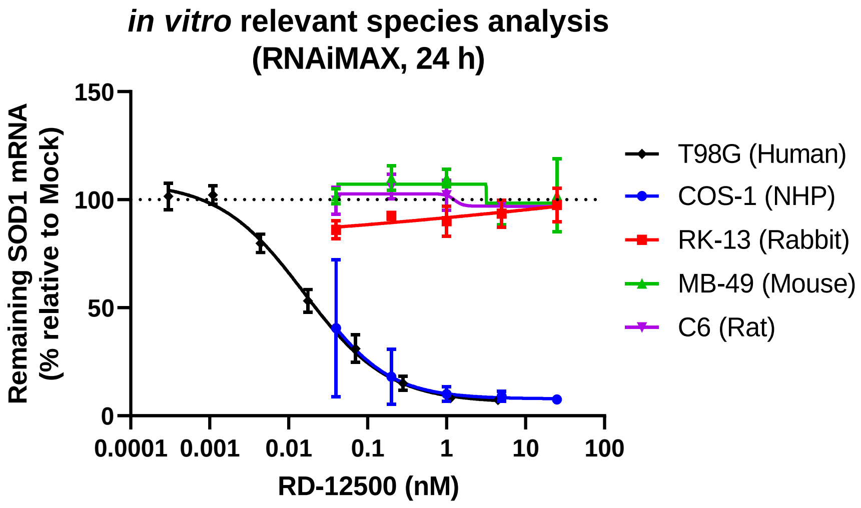


**Supplementary Figure 5. *In vitro* cross-species activity of RAG-17.** *In silico* analysis provided mismatch information between different species-derived SOD1 transcripts and "seed" region of RD-12500 guide strand. Species specific cell lines was transfected with RD-12500 at multiple concentration for 24 h and Species specific SOD1 mRNA expression was assayed by RT-qPCR.

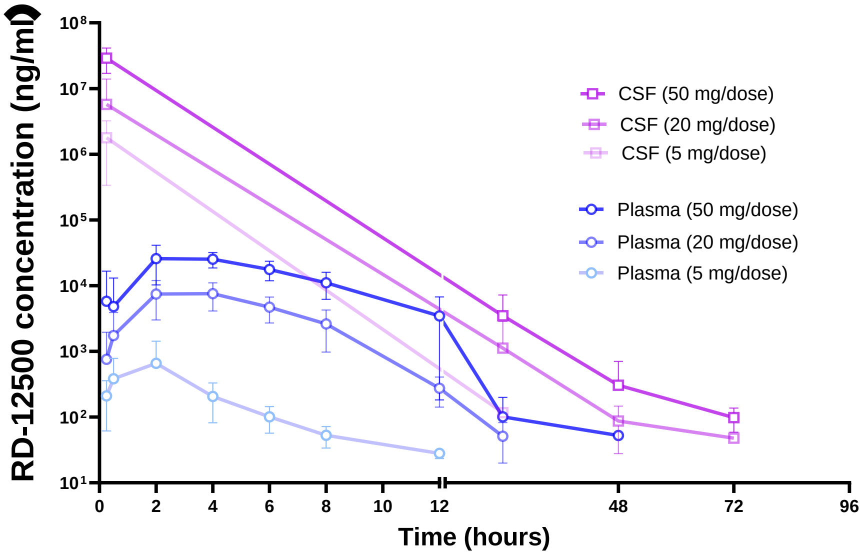


**Supplementary Figure 6. Pharmacokinetics of RD-12500 in CSF and plasma.** CSF and plasma were collected from cynomologus monkeys IT dosed with RD-12500 5, 20 and 50 mg/dose at time points including predose,15 min, 24 h, 48 h and 72 h (only for CSF) postdose and analyzed by validated LC-MS/MS methods. Each group included 10 animals (5 males and 5 females). Data represents mean ± SD.

**Supplementary Table 1.** ***in silico*** **analysis with perfect "seed" region complementarity to RD-12500 guide strand sequence.**

<table>
<tr><th>Gene</th><th>Off-Target Site Alignment</th><th></th><th>Prediction Algorithm</th><th>Site Location</th><th>3' Flanking Mismatches</th></tr>
<tr><td rowspan="3">UBAP2L</td><td>3' ACGUCCCGUAGUAGUUAAAGCU 5'</td><td>GS</td><td rowspan="3">miRanda</td><td rowspan="3">CDR</td><td rowspan="3">2</td></tr>
<tr><td>     |||| |||| |||||||||</td><td></td></tr>
<tr><td>5' GCCAGGACAUCCUCAAUUUCGU 3'</td><td>OT</td></tr>
<tr><td rowspan="3">CYTH2</td><td>3' ACGUCCCGUAGUAGUUAAAGCU 5'</td><td>GS</td><td rowspan="3">miRanda</td><td rowspan="3">3'UTR</td><td rowspan="3">4</td></tr>
<tr><td>     |:| || ||:  |||||||</td><td></td></tr>
<tr><td>5' UGCGGCGCUUCG-GAAUUUCGG 3'</td><td>OT</td></tr>
<tr><td rowspan="3">NDUFC2</td><td>3' ACGUCCCGU-AGU-AGUUAAAGCU 5'</td><td>GS</td><td rowspan="3">miRanda</td><td rowspan="3">3'UTR</td><td rowspan="3">4</td></tr>
<tr><td>      || ||| | :  ||||||||</td><td></td></tr>
<tr><td>5' UGGAGUGCAGUGGCGCAAUUUCGG 3'</td><td>OT</td></tr>
<tr><td rowspan="3">DISP2</td><td>3' ACGUCCCGU-AGU-AGUUAAAGCU 5'</td><td>GS</td><td rowspan="3">miRanda</td><td rowspan="3">3'UTR</td><td rowspan="3">4</td></tr>
<tr><td>      || ||| | :  ||||||||</td><td></td></tr>
<tr><td>5' UGGAGUGCAGUGGCACAAUUUCGG 3'</td><td>OT</td></tr>
<tr><td rowspan="3">KNOP1</td><td>3' ACGUCCCGUAGUAGU-UAAAGCU 5'</td><td>GS</td><td rowspan="3">miRanda</td><td rowspan="3">3'UTR</td><td rowspan="3">3</td></tr>
<tr><td>     ||||||| | ||| ||||||</td><td></td></tr>
<tr><td>5' GGCAGGGCAGC-UCAUAUUUCGG 3'</td><td>OT</td></tr>
</table>

GS, guide strand; OT, off-target site; CDR, coding region; 3'UTR, 3' untranslated region; G:U wobble base pairing is represented as ':'

**Supplementary Table 2. Tissue PK parameter of RD-12500.**

| Tissue | Male rats (n=3) | | | | Female rats (n=3) | | | |
|---|---|---|---|---|---|---|---|---|
| | $C_{max}$ (ng/g) | $T_{max}$ (h) | $AUC_{0\text{-}last}$ ( ng•h/g ) | $T_{1/2}$ (days) | $C_{max}$ (ng/g) | $T_{max}$ (h) | $AUC_{0\text{-}last}$ (ng•h/g ) | $T_{1/2}$ (days) |
| Liver | 18200 | 6.00 | 3640000 | 6.2 | 29900 | 6.00 | 4140000 | 8.1 |
| Kidney | 55800 | 6.00 | 11300000 | 6.6 | 75600 | 6.00 | 33100000 | 9.2 |
| Lung | 677 | 6.00 | 172000 | 21.9 | 636 | 144 | 234000 | NR |
| Heart | 521 | 6.00 | 191000 | 25.0 | 722 | 6.00 | 222000 | 8.6 |
| Muscle | ND | ND | ND | ND | ND | ND | ND | ND |
| Spleen | 7910 | 6.00 | 3370000 | 13.0 | 11100 | 6.00 | 2910000 | 11.5 |
| Brain | 3230 | 144 | 2340000 | NR | 2900 | 6.00 | 1310000 | NR |
| Brain stem | 3510 | 144 | 2160000 | NR | 2950 | 6.00 | 1040000 | NR |
| Cerebellum | 7410 | 6.00 | 4470000 | NR | 8370 | 6.00 | 2160000 | NR |
| Cervical spinal | 7770 | 6.00 | 4020000 | 27.3 | 6040 | 6.00 | 1850000 | NR |
| Frontal cortex | 1670 | 144 | 1400000 | NR | 1790 | 6.00 | 858000 | NR |
| Hippo | 2410 | 144 | 1400000 | NR | 3180 | 6.00 | 802000 | NR |
| Lumber spinal | 65900 | 6.00 | 22600000 | 21.0 | 75000 | 6.00 | 13500000 | NR |
| Striatum | 304 | 144 | 197000 | 20.1 | ND | ND | ND | ND |
| Thoracic spinal | 26100 | 6.00 | 11800000 | 23.1 | 26500 | 6.00 | 6480000 | NR |

"ND": not determined as more than half (>50%) of the individual values are not quantifiable. "NR": not reportable; "Brain": the remaining brain tissues.

**Supplementary Table 3. Primer sequences.**

| Genes | Species | Forward primer (5'-3') | Reverse primer (5'-3') |
|---|---|---|---|
| SOD1 | Human | aagcattaaaggactgactgaagg | caagtctccaacatgcctctc |
| CYTH2 | Human | tactttgagtacaccacggacaa | cctctcggatgctcagattctc |
| NDUFC2 | Human | ccttacggtttctgccggat | ctggcgatgcaaaccagc |
| DISP2 | Human | ctgcccacttcacctatccc | ggctcgttcctgcctgac |
| KNOP1 | Human | cgcacatagaccaggtgagg | agcttccgttttgcctgact |
| UBAP2L | Human | gtcgcctggatcttcagaca | acagccagagcaggaatctta |
| TBP | Human | tgctcacccaccaacaatttag | tctgctctgactttagcacctg |
| Sod1 | Macaca fascicularis | aaggtgtggggaagcattac | ccaccgtgttgtctggatag |
| Gapdh | Macaca fascicularis | atcaccatcttccaggagcga | ttctccgtggtggtgaagacg |
| Sod1 | Rabbit | gggacgcataacaggactga | tgcactcgtacagccttgtg |
| Tbp | Rabbit | tgaagttccccatacggctg | ctgctctgactttagcacctgt |
| Sod1 | Rat | agcggatgaagagaggcatgt | cttcatttccacctttgccca |
| Gapdh | Rat | atcaccatcttccaggagcga | ttctccatggtggtgaagacg |
| Sod1 | Mouse | ttggccgtacaatggtggtc | acagtttaatggtttgagggtagc |
| Gapdh | Mouse | aactttggcattgtggaagg | ggatgcagggatgatgttct |
